# Bio-inspired neural networks implement different recurrent visual processing strategies than task-trained ones do

**DOI:** 10.1101/2022.03.07.483196

**Authors:** Grace W. Lindsay, Thomas D. Mrsic-Flogel, Maneesh Sahani

## Abstract

Behavioral studies suggest that recurrence in the visual system is important for processing degraded stimuli. There are two broad anatomical forms this recurrence can take, lateral or feedback, each with different assumed functions. Here we add four different kinds of recurrence—two of each anatomical form—to a feedforward convolutional neural network and find all forms capable of increasing the ability of the network to classify noisy digit images. Specifically, we take inspiration from findings in biology by adding predictive feedback and lateral surround suppression. To compare these forms of recurrence to anatomically-matched counterparts we also train feedback and lateral connections directly to classify degraded images. Counter-intuitively, we find that the anatomy of the recurrence is not related to its function: both forms of task-trained recurrence change neural activity and behavior similarly to each other and differently from their bio-inspired anatomical counterparts. By using several analysis tools frequently applied to neural data, we identified the distinct strategies used by the predictive versus task-trained networks. Specifically, predictive feedback de-noises the representation of noisy images at the first layer of the network and decreases its dimensionality, leading to an expected increase in classification performance. Surprisingly, in the task-trained networks, representations are not de-noised over time at the first layer (in fact, they become ‘noiser’ and dimensionality increases) yet these dynamics do lead to de-noising at later layers. The analyses used here can be applied to real neural recordings to identify the strategies at play in the brain. Our analysis of an fMRI dataset weakly supports the predictive feedback model but points to a need for higher-resolution cross-regional data to understand recurrent visual processing..

## 1 Introduction

Performance on challenging visual tasks can increase when more processing time is allowed. Backward masking studies—wherein a meaningless stimulus meant to interrupt neural processing is shown with a varying delay after a target stimulus—have in particular demonstrated the importance of uninterrupted visual dynamics, even in response to static images (Kahneman, 1968). Such studies have also indicated that simple high-level visual categorization tasks can be achieved with very little processing time, for example on images presented for as few as 20ms, if the image is not degraded (Fabre-Thorpe, 2011; Thorpe, Fize, & Marlot, 1996). This suggests that a feedforward pass through the visual hierarchy is sufficient for these tasks. Noisy or degraded images, however, are classified with significantly lower accuracy with such brief processing times, but allowing more processing time before the onset of the mask increases performance on these more challenging images (Benoni, Harari, & Ullman, 2020; Johnson & Olshausen, 2005; Tang et al., 2018; Wyatte, Curran, & O’Reilly, 2012). Neural investigations have indicated a role for recurrent connections in influencing neural dynamics during masking experiments (Lamme, Zipser, & Spekreijse, 2002; Lamme et al., 2002; Rajaei, Mohsenzadeh, Ebrahimpour, & Khaligh-Razavi, 2019; Wyatte, Jilk, & O’Reilly, 2014). Therefore, recurrence is believed to be important for making sense of visual inputs under imperfect conditions.

The ability of recurrence to increase performance on static image classification tasks is perhaps surprising. Naively, we may expect that simply re-processing the same information over and over through recurrent connections would only make the visual system more confident in its initial (incorrect) classification; yet it instead allows it to produce a different output. Many different descriptions of the computational functions that recurrence implements that make it so useful have been put forth based on both functional and anatomical data. Feedback recurrence, wherein a later visual area sends connections back to an earlier one, has been implicated in figure-ground segregation, high level perceptual grouping, prediction, and ‘explaining away’ (Hupé et al., 1998; Kim, Linsley, Thakkar, & Serre, 2019; Kreiman & Serre, 2020; Mumford, 1992; Murray, Schrater, & Kersten, 2004). Connections within a visual area—known as horizontal or lateral connections—are associated with more low level functions such as spatial sharpening, cross-feature competition and normalization, and small scale visual grouping (Jones, Grieve, Wang, & Sillito, 2001; Kim et al., 2019; Stemmler, Usher, & Niebur, 1995; Stettler, Das, Bennett, & Gilbert, 2002).

Convolutional neural networks (CNNs) are artificial neural networks inspired by the structure of the visual system and frequently used as models of visual processing in the brain (Lindsay, 2021; Yamins & DiCarlo, 2016). While many of the original studies using CNNs focused on feedforward networks only, studies are now exploring the role of recurrence using these models. Networks trained to perform tasks with recurrence have been used to show that recurrence makes these networks better at a variety of challenging visual problems (Kang & Druckmann, 2020; Kim et al., 2019; Linsley, Kim, Veerabadran, Windolf, & Serre, 2018; Spoerer, McClure, & Kriegeskorte, 2017) and better at predicting neural data (Kar, Kubilius, Schmidt, Issa, & DiCarlo, 2019; Kubilius et al., 2019; Nayebi et al., 2021). This suggests they are useful models for probing how recurrence functions in the visual system.

Here, we compare different ways of adding recurrence to a CNN in order to understand the functions recurrence may play in enhancing classification of degraded images. Specifically we start with a feedforward network trained to classify noise-less digits and add reccurence to this pre-trained network to help the network classify noisy images. This procedure of starting with a fixed feedforward network has three advantages. The first is that it ensures that the feedforward pass will be able to classify clean images, thus replicating behavioral results mentioned above. Second, it makes it possible to directly compare the impact of different types of recurrence on the same neurons in the same network. Third, there is experimental evidence, particularly with respect to feedback connections, indicating that recurrent connections mature after feedforward ones developmentally (Batardière et al., 2002; Markov & Kennedy, 2013). This suggests recurrent connections may be refined within the context of an already matured feedforward pathway.

We try out different ways of adding both feedback and lateral connections. First, we use bio-inspired methods: we add lateral connections that implement surround suppression, which is found throughout the visual system (Angelucci et al., 2017; Hasani, Soleymani, & Aghajan, 2019); for feedback connections, we implement a modified predictive processing scheme (Choksi et al., 2020, 2021; Mumford, 1992; Rao & Ballard, 1999). To add connections inspired mainly by the function we know recurrence to play, we also add lateral and feedback connections trained with backpropagation to classify noisy images.

We compare the impact these four different ways of adding recurrence have on the classification of noisy and degraded images. While most studies of recurrence in CNNs have focused on showing what kind of training and architectures lead to the best performance, we instead aim to understand how these different forms of recurrence work at the neural and behavioral level. This is achieved through both exploratory and hypothesis-driven analyses of network behavior and neural activity.

## 2 Results

We first train a feedforward CNN, which we refer to as the feedforward core, on the standard MNIST handwritten digit recognition task (see Methods 4.1). Starting from the same feedforward core, we then build four different models, each of which has recurrence added in a different way.

For our analyses, we report results from five feedforward cores trained starting from different random initializations. For each core, we train two instantiations of each of the types of recurrent connections (with the exception of surround suppression which has no trainable parameters).

### 2.1 Different ways of adding recurrence all increase performance on challenging images over time

The feedforward core we use (Figure 1A, see Methods 4.1) has 4 layers in total: two convolutional layers (each with a nonlinearity and max-pooling), a fully-connected layer, and an output layer (with one unit for each of the 10 digit classes). To add lateral recurrence to this network, we add convolutional recurrent connections at the first convolutional layer (‘layer 1’) (Figure 1D). Feedback recurrence is added from the second convolutional layer (‘layer 2’) to the first (Figure 1C). These connections must learn to work within the constraints of the existing feedforward core, which is not updated in light of the addition of recurrence. The addition of recurrence to a feedforward network creates temporal dynamics.

**Figure 1:**
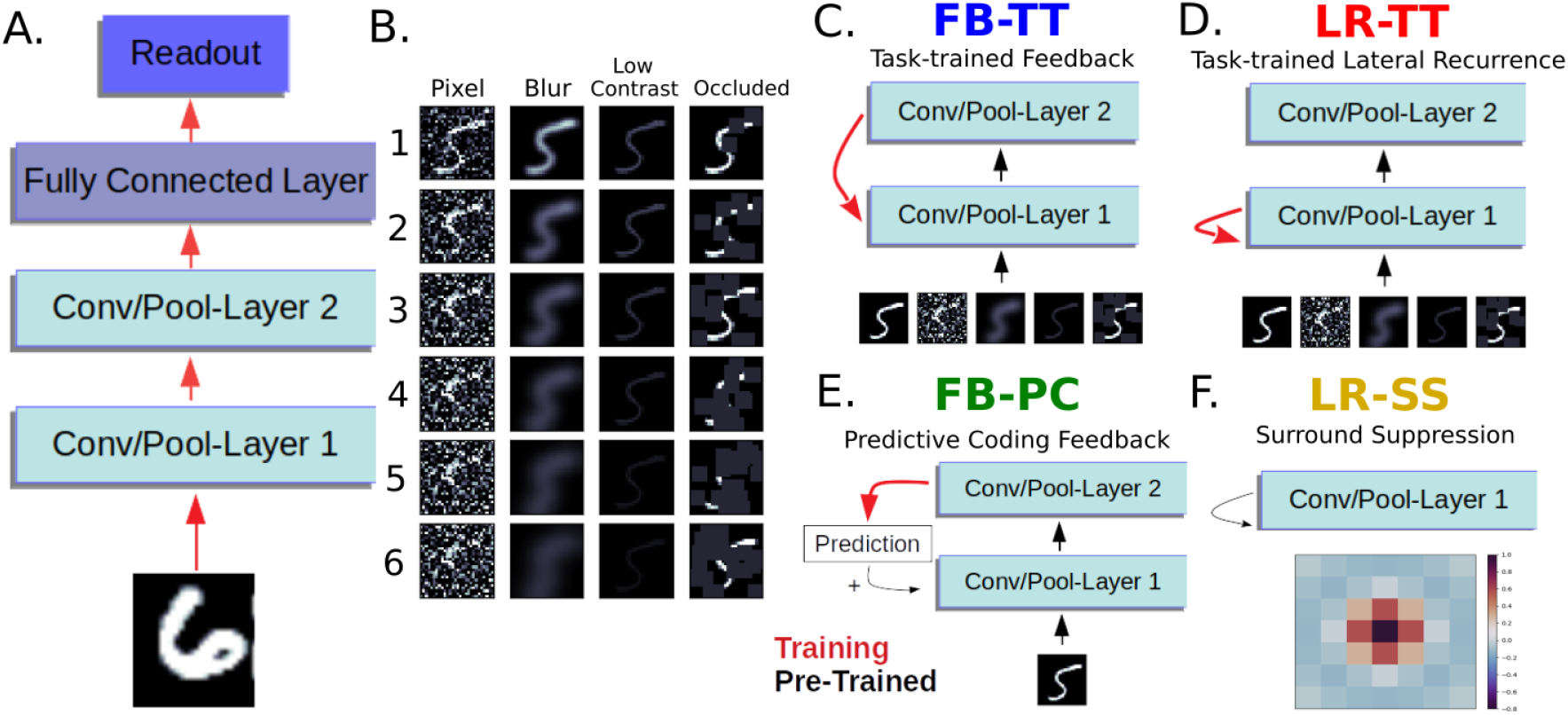
Network architectures and training data. A. Architecture of the feedforward core. The first convolutional layer has 32 feature maps, the second has 64, and the fully connected layer has 1024 units (more details in Methods 4.1) The feedforward core is trained on clean images only. B. Different image degradations used. Columns indicate noise type and rows levels. Analyses were performed on noise level 5 (task-trained networks were trained on level 3). Note: low contrast images are plotted on a different colorscale as even level 1 is too faint to see on the normal scale. C-F. Four different forms of recurrence are added to the pre-trained feedforward core. C. Task-trained feedback is convolutional, added from the second convolutional layer to the first, and trained using clean and noisy images. D. Task-trained lateral connections are convolutional, added amongst units in the first convolutional layer, and trained using clean and noisy images. E. Predictive feedback connections are from the second layer are convolutional and trained to predict layer 1 activity in response to clean images. These predictions are added to layer 1 activity. F. Convolutional spatial surround suppression (within a single feature map) is added to the first layer using the filter shown. This network does not involve training.

We explored biologically-inspired options for adding feedback and lateral connections (see Methods 4.1). For feedback we implement a predictive processing scheme introduced in Choksi et al. (2020) wherein feedback connections are trained to re-construct layer 1 activity from layer 2 activity (in response to clean images only) (Figure 1E). At run time, this reconstructed activity is then added to the true activity at the first layer. While this form of predictive processing differs from the more common implementation of predictive coding (Rao & Ballard, 1999) in that it does not propagate error signals, it has been shown to increase performance on classification of images with pixel noise (Choksi et al., 2020, 2021). For the lateral connections, we implement within-feature spatial surround suppression and nearby facilitation by applying a one channel convolutional filter defined by a difference of Gaussians (Figure 1F), inspired by Hasani et al. (2019) (though note in their study the full model is trained with this filter present, which is not the case here).

We compare these models to others with matching anatomy of recurrence (i.e. feedback or lateral) but with weights trained explicitly to help the network classify degraded images. Specifically, in our ‘task-trained’ networks, the added recurrent connections alone are trained through backpropagation to classify images with four different types of level 3 noise added: pixel noise, blur, low contrast, and occlusion (Figure 1B). The network is run for three timesteps, which results in two applications of recurrent influence after the feedforward pass. Training loss is calculated at the final timestep.

After creating these four networks, we then ask if they can each replicate the basic findings from backward masking experiments. Specifically, we want the models to (a) perform well on clean images with only one time step (that is, after just the feedforward pass), (b) show worse feedforward performance with increasing noise, and (c) show an increase in performance on noisy images with increasing time. Because the feedforward core was trained to classify clean images only, both (a) and (b) are captured by all four networks by default.

We find that all four networks also capture (c). In Figure 2A, we show performance as a function of noise level and time (averaged over all noise types). As expected, the task-trained networks outperform the bio-inspired ones as they are directly optimized for the task (though note task-trained networks were trained only on noise level 3). Surround suppression also outperforms predictive feedback. All recurrent networks show a significant (*p* < .05) increase in performance compared to the feedforward network (timestep 0) for noise levels 2-5 (surround suppression is not significant at level 1 and predictive feedback is not at level 6). The rest of the analyses will use images with noise level 5.

**Figure 2:**
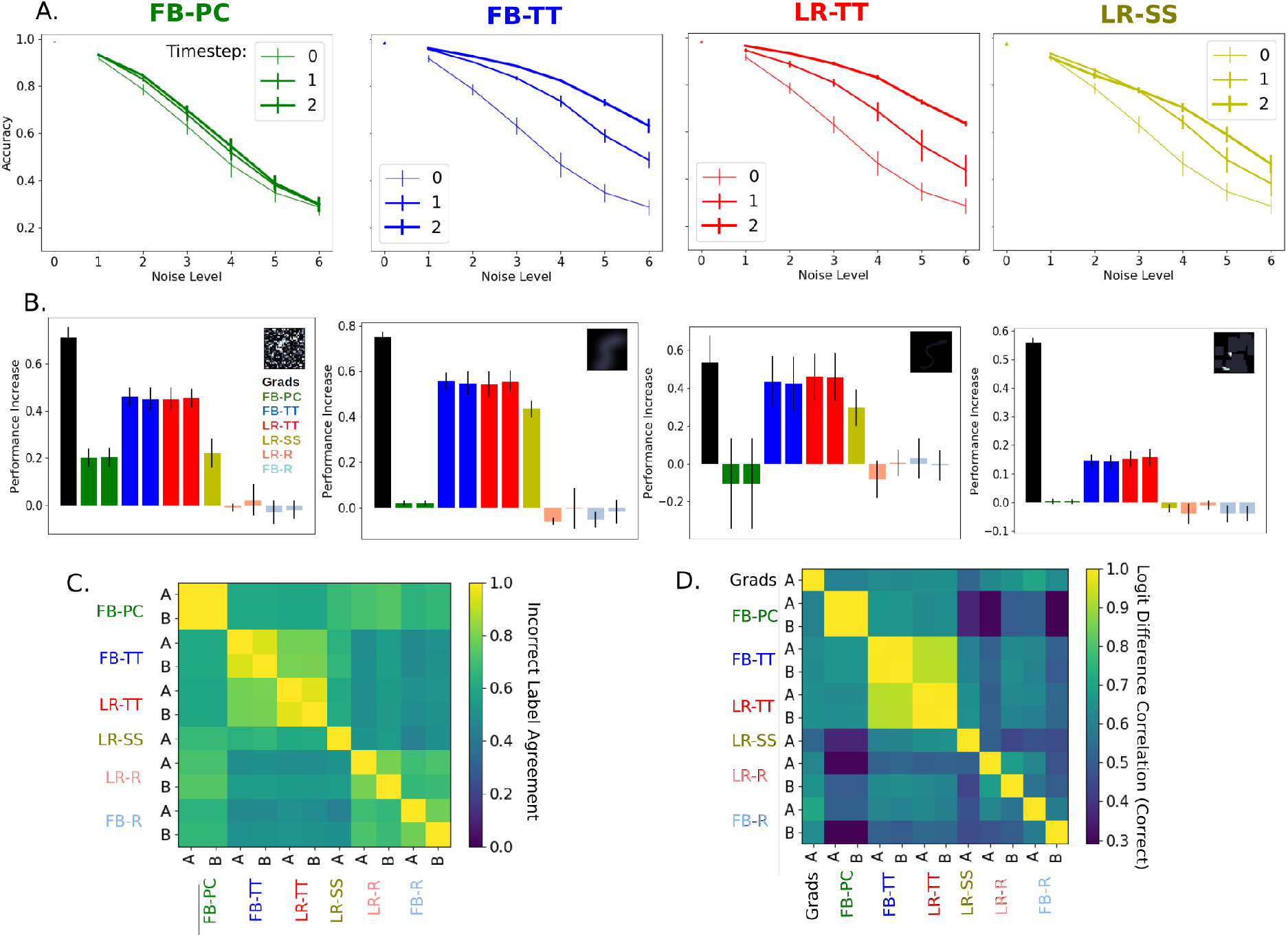
Classification behavior across different networks. A. Performance of different networks on different noise levels over time (averaged over all noise types). Timestep 0 is the feedforward performance and is thus the same for all networks. B. Performance increase (performance with recurrence minus feedforward performance) for each recurrent network type broken down by noise type (at level 5). Performance is shown separately for the two instantiations of each recurrent option trained (with the exception of surround suppression which has no trainable parameters). In addition, the performance increase that comes from applying a gradient-based update to first layer activity is shown (‘Grads’), as is performance with random recurrence (‘LR-R’ and ‘FB-R’). All errorbars are ± s.t.d. over feedforward cores. C. Incorrect label agreement across networks is defined as: of the set of images that both networks classify incorrectly, the fraction of images that are given the same label by both networks. D. For correctly classified images, we report the correlation between the last layer activity (the ‘logits’) at the final timestep (with activity at the first timestep subtracted). Networks labeled A and B refer to the two different instantiations of recurrence trained. Matrices are the mean over all feedforward cores.

Note: networks trained to classify both clean and noisy images with recurrence from the start do not replicate (a); that is, they need multiple time steps to reach peak performance on clean images (not shown; a similar effect has been observed elsewhere (Rajaei et al., 2019).). This makes them inappropriate for studying the phenomena observed in masking studies.

### 2.2 Classification patterns differ between task-trained and bio-inspired networks

To investigate if these different forms of recurrence work the same way, we first analyze classification performance in more detail.

Different types of recurrence may be more or less suited for handling different types of noise; for example, surround suppression sharpens image representations, making it well-suited to counter blur. In Figure 2B, we break performance increases down by the four different types of noise added. We also compare performance increases to those attained by applying a large gradient step to the first layer activity, calculated on an image-by-image basis to increase classification accuracy.

Across the two instantiations of each kind of recurrence we see strong agreement in how much recurrence increases accuracy. The addition of random (untrained) recurrence does not lead to performance increases.

Despite the fact that task-trained feedback and predictive feedback use similar anatomy, they differ in their patterns of performance across noise types. Specifically, predictive feedback only significantly increases performance on images with Gaussian pixel noise (which was shown in Choksi et al. (2020)) and blur. Task-trained feedback, on the other hand, performs well on all noise types and performs best (relative to performance enhancement from gradients) on low contrast noise. Task-trained lateral recurrence has a nearly identical pattern of performance as task-trained feedback and differs from that of surround suppression. Surround suppression performs best on blur but also significantly increases performance for images with pixel noise and low contrast. These results suggest that training is more relevant for the function of recurrence than anatomy in these networks.

Comparing the mistakes made by the different networks can also offer insights into their respective behavior. In Figure 2C we show the number of images wherein both networks came to the same incorrect conclusions, as a fraction of the total number of images they both classified incorrectly. Like the accuracy measures, this shows large agreement between the two different instantiations of the same recurrence type (blocks on the diagonal for FB-PC, FB-TT, and LR-TT). It also shows agreement between the two task-trained networks (large block for FB-TT and LR-TT).

There is also some agreement according to this metric amongst the networks that receive random recurrence. This is inherited from the errors the feedforward core makes and therefore doesn’t represent similarities between these types of recurrence. To compare classification behavior without this confound, we look at the activity of the 10-unit final layer (known as the logits). Specifically, we subtract the logit activity at the first timestep from that at the final timestep to isolate the influence of recurrence. We also look here only at images correctly classified by both networks. These logit differences thus indicate how confidence across the different digit identities is changed through recurrence, but is not influenced simply by the fact that task-trained networks perform better. As we show in Figure 2D, the correlation between this measure is highest for different instantiations of the same recurrence but closely followed by the correlation between the task-trained forms of recurrence.

These comparisons show that the task-trained networks behave more similarly to each other than they do to their bio-inspired counterparts with the same anatomy of recurrence.

### 2.3 Task-trained recurrence influences neural activity differently than bioinspired recurrence

In addition to comparisons at the output or ‘behavioral’ level, we can also compare how each of the different forms of recurrence impacts layer 1 activity. Because all of the recurrent models share the same feedforward core, these comparisons can be done directly.

To get a broad sense of the influence that recurrence has on layer 1 activity we plot histograms of the ‘recurrent influence’. This is simply the difference between activity at the final timestep and that at the first timestep. In Figure 3C, we show example histograms from individual networks. All networks have a slightly positive mean influence but show both increases and deceases in activity. Predictive feedback has a smaller mean than the other networks and surround suppression has a much larger one (due mainly to its long positive tail).

**Figure 3:**
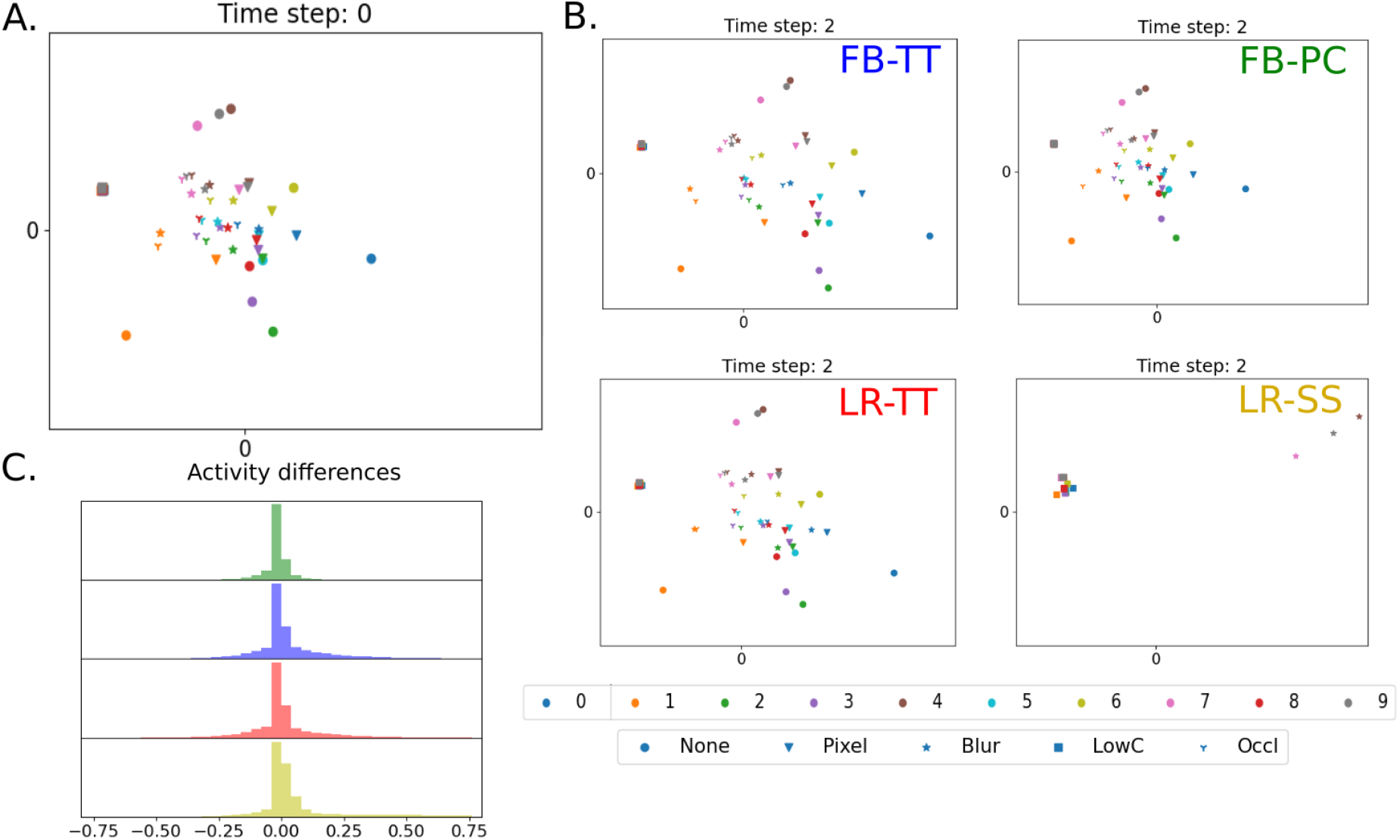
How recurrence impacts activity. A. Representation of digit and noise information in layer 1 activity in the feedforward network. Principal components are created using feedforward activity in response to clean images only (so as to capture the representational space of the feedforward core) and the feedforward activity in response to images of all noise types is shown (indicated by marker type; digit identity is indicated by color). Marker location signifies mean across all images of that noise and digit identity. B. The same space as in A but showing the representation after the influence of each type of recurrence. C. Histograms of recurrent influence (the change in activity that comes from recurrence). Histograms are made from all units in the first layer in response to images of all noise types, from example networks. Mean and standard deviation of mean differences over all feedforward cores: FB-TT .19±.12, LR-TT .10±.09, FB-PC .02±.03, LR-SS .94±.38.

We can also get a rough impression of the impact of different forms of recurrence by visualizing how layer 1 activity changes over time. Specifically, we are interested in how recurrence impacts image representation as a function of digit identity as well as image noise. To perform this visualization we first reduce layer activity down to two dimensions. To do this, we perform principal components analysis (PCA) on layer 1 activity at timestep 0 in response to clean images only. This captures the variance relevant to how the feedforward core was trained and may therefore highlight changes that impact the downstream classifier. As all models share the same core, this also allows for direct comparisons of their respective visualizations. In Figure 3A we show first the representation of images at time step 0 for an example feedforward core. We then show the representation at the final timestep for the recurrent networks (Figure 3B).

These plots highlight several features of the representation. First, in the feedforward response we can see clean images of different digits (circles) spread out in a roughly circular shape. The noisy variants of these images are closer to the center and thus less separable. With recurrence, these patterns spread out, particularly for task-trained networks and for surround suppression (indeed, the activity corresponding to most noise types does not remain in this frame for the surround suppression network) The influence of predictive feedback, on the other hand, does not expand activity in this space; rather, it rearranges the representations of noisy images within the existing feedforward activity area.

To more directly compare the impact of recurrence in these different networks, we plot the correlation between the recurrent influence from different networks averaged over a set of images containing all noise types. This indicates the extent to which the different forms of recurrence push layer 1 activity in the same way in response to the same image. As can be seen in Figure 4A, the task-trained networks are strongly correlated with each other, far more than they are with their bio-inspired counterparts. Furthermore, these task-trained networks are also not correlated with the activity changes calculated by backpropagation (that is, the gradients). This suggests that the similarities between the task-trained networks are not simply the result of them both following the gradient signal that they were trained on.

**Figure 4:**
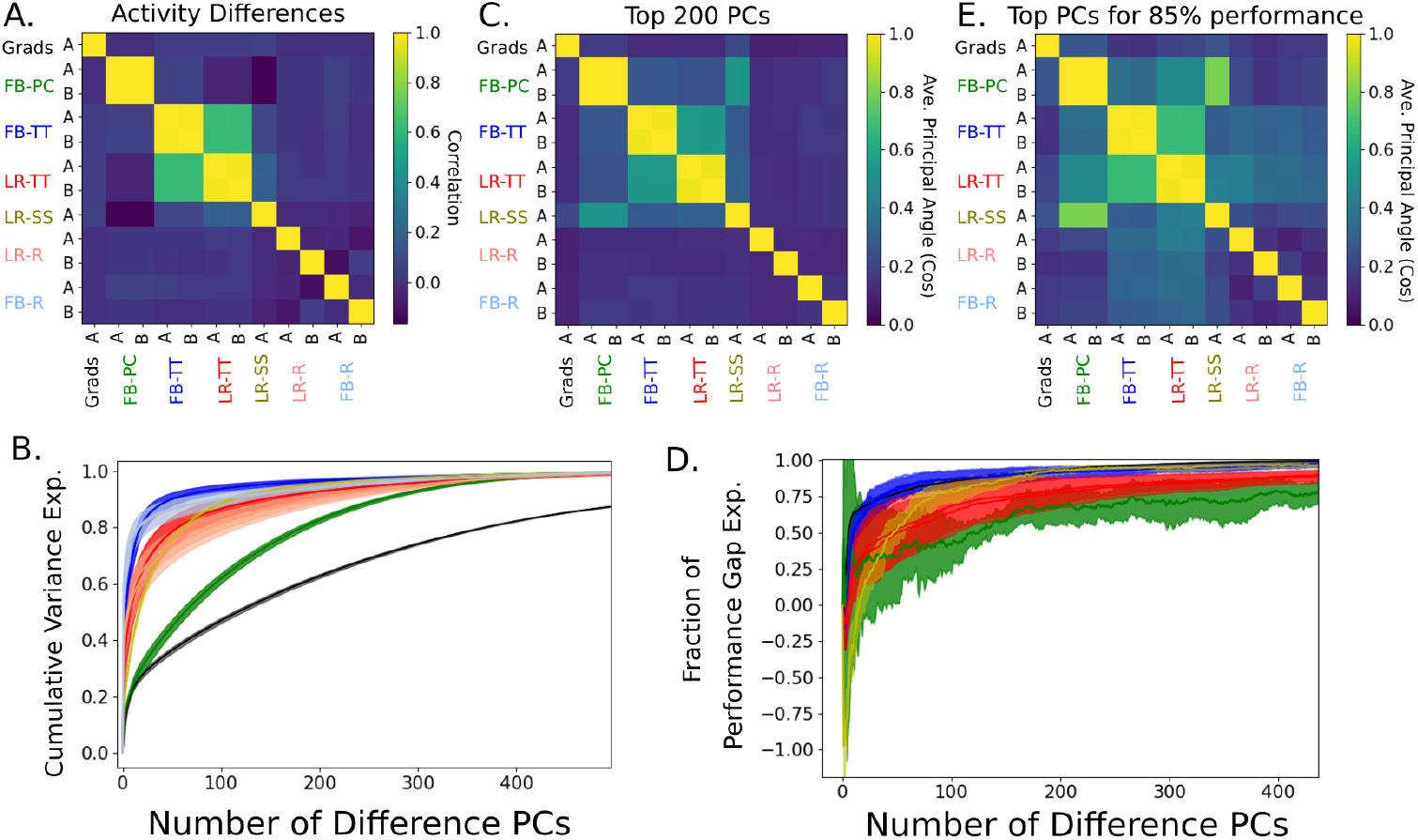
Comparing Recurrent Influences. A. The correlation coefficient of recurrent influences (defined as activity after recurrence minus feedforward activity) across networks in response to the same set of images (of all noise types). B. Cumulative variance explained by the top 500 ‘difference’ PCs (PCs made from the recurrent influences); colors are is in previous figures. C. Subspace alignment (as measured by the cosine of the average principle angles) according to the top 200 difference PCs. D. How much of the performance gap (defined as performance with recurrence minus feedforward performance) is achieved when recurrent influences are recreated using the indicated number of difference PCs. E. Subspace alignment when using the number of PCs needed to reach 85% of the performance gap for each network (Grads: 95, FB-PC: 666, FB-TT: 71, LR-TT: 306, LR-SS: 141). Shaded regions are ± s.t.d. over all feedforward cores. Matrices are means over all feedforward cores.

It is possible that differences on the individual neuron level are still consistent with the implementation of a similar larger-scale influence. To explore the overall patterns in how recurrence impacts layer 1 activity, we perform PCA on the recurrent influences, creating what we refer to as ‘difference’ PCs. The cumulative variance explained by these PCs is shown in Figure 4B. This shows the dimensionality of the influence. Interestingly, this dimensionality appears to relate somewhat to the anatomy of the recurrence. Lateral recurrence specifically (including task-trained, surround suppression, and random) require a similar number of PCs to reach a given amount of variance, whereas task-trained and random feedback require fewer. Predictive feedback however is higher dimensional by this measure.

To determine if the dominant patterns of recurrent influence are similar across the different networks, we measure the extent to which the subspaces defined by the first 200 PCs of each network are aligned. To measure alignment we take the cosine of the average of the principal angles between the two subspaces. According to this measure, 0 is unaligned and 1 is aligned. As can be seen in Figure 4C, the subspaces of the task-trained networks are more aligned with each other than with the bio-inspired networks.

These difference PCs are calculated based on the amount of variance they explain in the neural activity changes. This does not guarantee that these patterns of activity changes are actually responsible for the increased performance observed with recurrence. To address this, we reconstruct the recurrent influence using a subset of the PCs and see how well this increases performance (Figure 4D; see Methods 4.3). We then identify for each network the set of PCs that captures 85% of the performance increase that comes with recurrence. Calculating subspace alignment using these PCs still shows again that task-trained networks are more aligned with each other than with bio-inspired networks. Interestingly, both of these alignment measures find some similarity between the two bio-inspired networks; this would suggest that perhaps some of the activity changes used by predictive feedback and surround suppression to enhance performance are similar.

In total, these analyses show that the changes in neural activity elicited by recurrence in the first layer are similar for task-trained networks. These changes differ from those induced by bio-inspired recurrence; the extent to which the bio-inspired methods differ from each other depends on the metric used.

### 2.4 Task-trained networks de-noise image representations at later but not earlier layers

Having established that task-trained networks work differently than bio-inspired ones, we now set out to understand how these networks enhance performance on challenging images.

One natural hypothesis regarding the role of recurrence in the visual system is that of ‘de-noising’. That is, recurrent influences may push the representation of a noisy image to be more like the representation of the same image without noise, thus allowing the system to classify the image as though it were noiseless. Here, we present the network with the same digit image with and without the four different types of noise added. We then measure the correlation between the feedforward network activity in response to the clean and to the noisy image (see Methods 4.3). The extent to which this correlation increases over time is a measure of how much the activity has been de-noised by recurrence.

In Figure 5, we show the amount of de-noising in each network broken down by noise type and layer (calculated specifically on images that went from being incorrectly to correctly classified with recurrence). Looking at the predictive feedback network, we see that it does in fact do de-noising; that is, the predictive feedback makes the representation of the noisy images at the first layer look more like that of their clean counterpart and this effect propagates to all layers in the network (this is consistent with similar results from Choksi et al. (2021)). Surround suppression shows a mixed effect, with some noise types exhibiting de-noising at layer 1 and others not.

**Figure 5:**
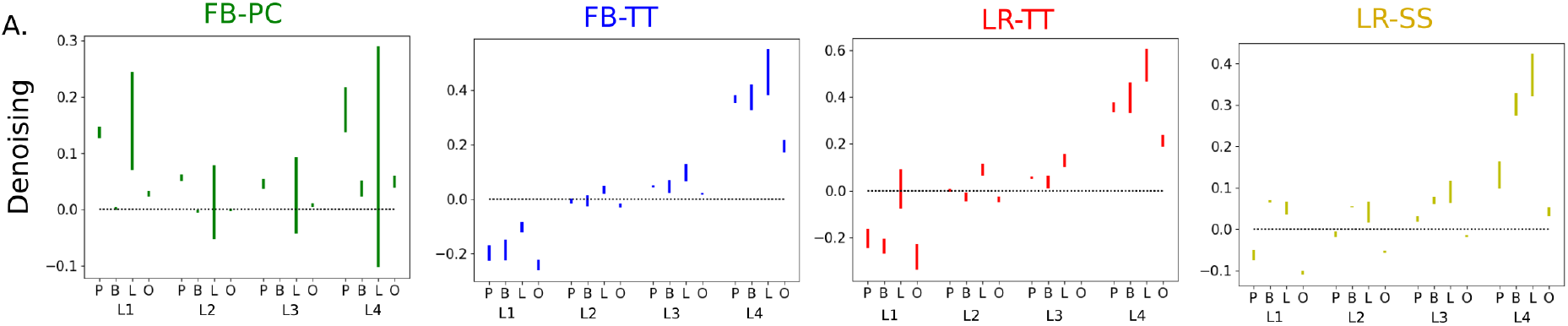
A. De-noising (the extent to which recurrence makes the representation of a noisy image more like that of its clean counterpart) for each network type broken down by noise type and layer. Images used were those that went from being incorrectly to correctly classified with recurrence.

Surprisingly, in the task-trained networks the first two layers do not show signs of de-noising. In fact, at the first layer recurrence predominantly makes the representation of the noisy image *less* like that of the clean one. At later layers, however, de-noising does occur. These dynamics at later layers are solely the result of changes at layer 1; therefore, the moving of noisy images away from their clean counterparts in layer 1 activity space counter-intuitively causes the opposite effect at layer 4. This indicates that the feedforward weights funnel activity in such a way that de-noising at the first layer is not the only means of enhancing performance via de-noising at the final layer. Rather, it seems the task-trained recurrence is able to explore different regions of layer 1 activity space than is normally used by the feedforward core in order to achieve this effect.

### 2.5 Predictive feedback and task-trained networks use different strategies to increase cross-layer dimensionality

To gain more intuition into exactly how the networks carry out this transformation across layers, we again plot activity according to noise and digit identity in a low-dimensional space (Figure 6). As opposed to Figure 3A and B, the principal components in this case are created from feedforward activity in response to noisy and clean images in order to capture variance related to noise. We can see, particularly at the first two layers, that initially images of different noise types form separate clusters, with digits arranged similarly within them. Task-trained recurrence and surround suppression spread these clusters out. At the third layer, surround suppression and task-trained recurrence help the representation shift to align according to digit identity along the y-axis and noise identity along the x-axis; this representation is much more conducive to classification by digit identity independent of noise. By the fourth layer activity is organized more by digit identity than by noise type.

**Figure 6:**
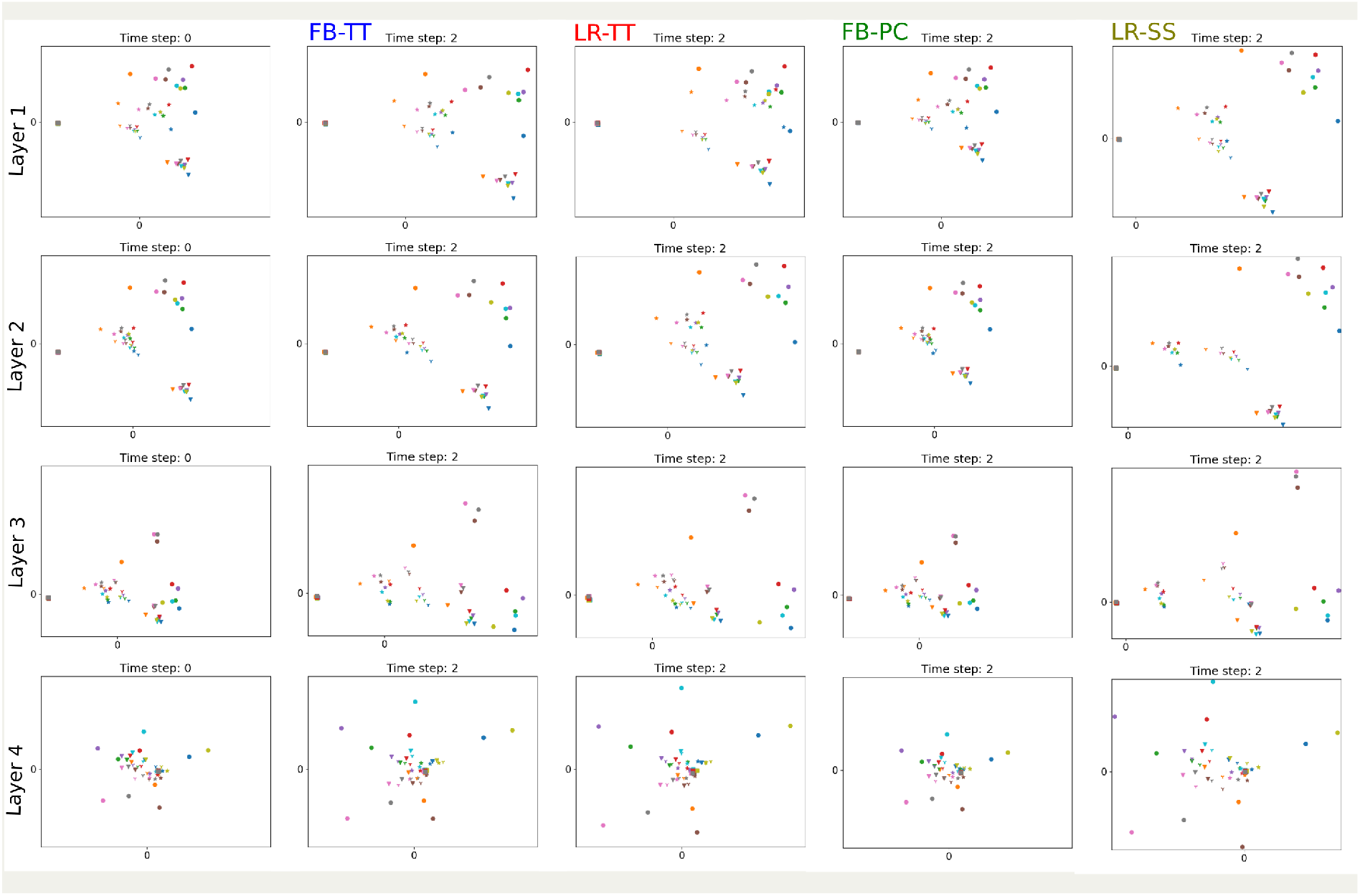
How recurrence influences representations at all layers, in example networks. PCs are made using feedforward activity in response to images of all noise types and clean images; therefore axes are the same for all plots within an indicated layer (though note the scale is different for the surround suppression network). Images are colored according to digit identity and marker type indicates noise type as in Figure 3.

As suggested by the de-noising analysis, the predictive feedback model appears to keep activity within the region defined by the feedforward response. The task-trained networks and surround suppression on the other hand are pushing activity away from this region and generally expanding it. In this way, the ’anti-de-noising’ found at the first layers of these networks likely does not represent an active push of noisy images away from their clean counterparts in particular, but rather the result of all points becoming farther from each other (such expansion within and across noise types can be see in Supplementary Figure 9).

How can expansion help classification? It is known that increased dimensionality of a neural representation can make classification easier (Fusi, Miller, & Rigotti, 2016). While in this case, increased dimensionality at the first layer of the network is not directly responsible for increased classification performance (as classification occurs at the fourth layer) it is possible that in an analogous way, increased dimensionality at the first layer may increase the ability of that layer’s activity to drive relevant activity patterns at the second layer.

To explore these questions we used two metrics: layer dimensionality and cross-layer dimensionality (see Methods 4.3). Layer dimensionality is simply an estimate of the dimensionality of activity at a given layer based on principal components analysis. Cross-layer dimensionality is an analogous measure meant to capture the patterns of co-variance across two layers. Looking at the impact of recurrence on layer dimensionality (Figure 7A), we can see that for task-trained and surround suppression networks, recurrence increases the dimensionality of the activity at the first layer. Increased dimensionality at the final layer is also associated with increased classification performance in these networks, given that the representation of images that go from being incorrectly to correctly classified show an increase in dimensionality while the reverse do not.

**Figure 7:**
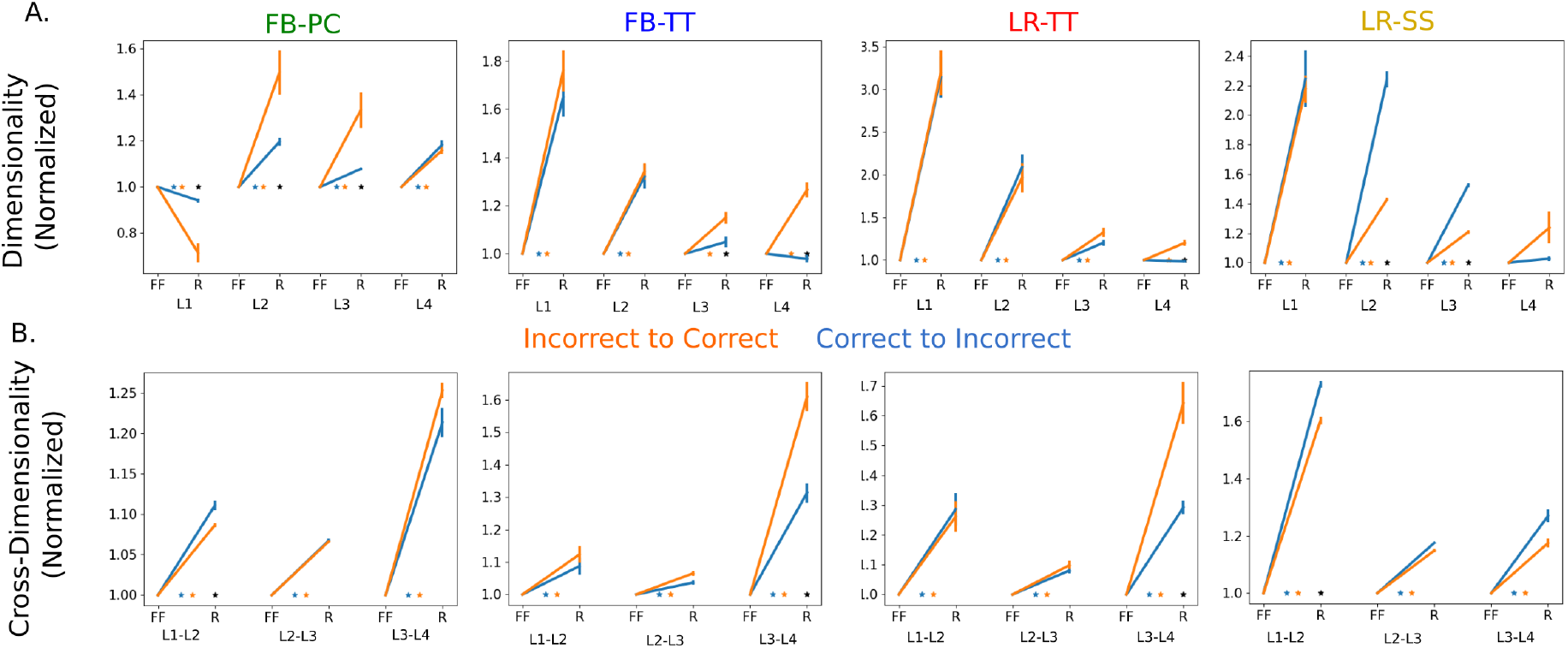
Impact of recurrence on dimensionality metrics. A. Layer dimensionality (normalized for each individual network by feedforward dimensionality) for all layers before (FF) and after (R) recurrence. B. Cross-layer dimensionality for indicated layer pairs (normalized for each individual network by feedforward cross-dimensionality) for all layers before (FF) and after (R) recurrence. Error bars are mean ± s.e.m. across all networks of each type. Color indicates classification behavior. Colored asterisks indicate significant (*p* < .01) change in dimensionality with recurrence; black asterisks indicate significant difference between two classification categories.

The predictive feedback network, however, shows a very different strategy not based on expanding dimensionality. Specifically, the dimensionality of activity at the first layer *decreases* as a result of the predictive feedback, particularly for images that go from being incorrectly to correctly classified. There is also not as strong a relationship between dimensionality at the last layer and performance.

Despite the dramatically different way that predictive feedback impacts dimensionality at layer 1, all forms of recurrence increase cross-layer dimensionality across layers 1 and 2 (and across other layer pairs; Figure 7B). This suggests that an important impact of recurrence may be to increase communication across layers in a hierarchy. In predictive feedback networks, this enhanced communication is achieved by making the noisy image representation more like the clean ones; in the other networks this is done by allowing activity to explore new dimensions of activity space.

### 2.6 Application of analyses to fMRI data

We demonstrated a difference between predictive feedback and other forms of recurrence on two metrics: denoising and layer dimensionality. Both of these metrics can be applied to real neural recordings as they only require a measure of neural activity over time in response to noisy and clean images. We analyzed an openly-available fMRI dataset wherein subjects were shown blurry and clean versions of the same images (see Methods 4.4) (Abdelhack & Kamitani, 2018). The results are shown for each of the 5 subjects individually in Figure 8. As can be seen in Figure 8A, no subject shows significantly negative de-noising values in any brain region in the visual system (in fact only 12 out of the 90 brain regions have a negative mean de-noising value at all). Two subjects (S03 and S04) show at least one brain region with significant positive de-noising values in both hemispheres, particularly at V1. This is in line with the predictions of how predictive feedback would influence this area. Predictive feedback is also expected to decrease dimensionality. Subject S04 shows strong decreases in dimensionality in line with the strong positive de-noising values across both hemispheres. Subject S03 shows one hemisphere with decreasing V1 dimensionality and one with increasing.

**Figure 8:**
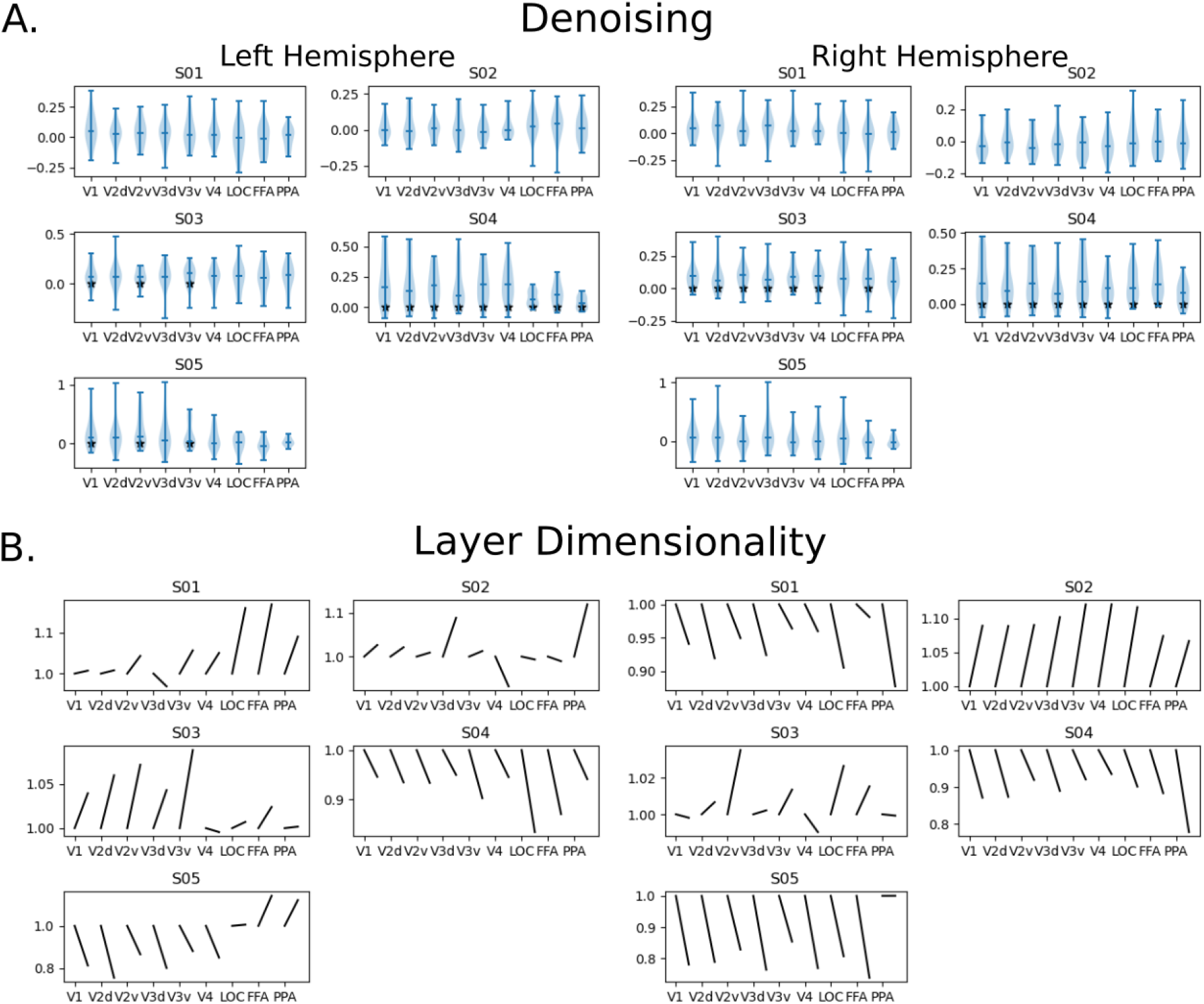
Analysis of fMRI data for 5 subjects. A. Distribution of de-noising values over all 20 image pairs (blurred and clean) for voxels from indicated brain regions. Lines indicate median values. Black asterisks indicate mean significantly (*p* < .01) different from zero. B. Normalized (by first time bin value) layer dimensionality of voxel activity calculated over all 40 images, shown for first and last time bins. Subject number indicated in titles; data broken down by hemispheres.

In total, one subject (S04) provides reasonably strong support for the predictive feedback model (though note that in our model dimensionality increased in later layers which it does not here; this could be due to predictive feedback occurring at all areas). No subjects provide strong support for the task-trained networks models.

Though not resolved in time, the analyses originally performed on this data in Abdelhack and Kamitani (2018) also generally support the notion that visual representations of blurred image are similar to that of their clean equivalents.

## 3 Discussion

Here we showed that many different styles of recurrence are able to enhance the ability of a feedforward network to classify degraded images, in line with behavioral findings. While it may be expected that recurrent connections with the same kind of anatomy—that is lateral vs. feedback connections—would function in a similar way, this is not what we find. Instead, both anatomical forms of recurrence function similarly when trained via backpropagation on the same task. These learned recurrent influences differ from those generated by lateral connections that implement surround suppression as well as from feedback connections that generate predictions.

This work conceptually replicates previous studies that showed the benefits of task-trained recurrence in CNNs, but shows that it also holds when the feedforward network is pre-trained and held fixed. Our work also explicitly replicates Choksi et al. (2020) which showed how this form of predictive feedback can enhance performance on Imagenet images with pixel noise and extends it to a new dataset and noise type (blur). Our result that post-hoc addition of surround suppression to a pre-trained CNN enhances performance is, as far as we know, a novel finding.

In some ways, the fact that the task-trained networks function similarly may not be surprising. They were both trained using gradient descent and would therefore be expected to learn to do what the gradient signals indicate are the best ways to change layer 1 activity. However, we know that task-trained recurrence doesn’t actually do a very good job of approximating the gradients (Figure 3A), which suggests this is not the best description of the strategy used by task-trained networks. The nature of recurrence requires the network to calculate its impact on layer 1 activity using only the activity of units at either layer 1 (lateral recurrence) or layer 2 (feedback). It therefore is surprising that both forms of recurrence appear to have found a similar strategy using these different neural populations.

This work has implications for how we use CNNs to study the visual system. It is common to compare networks with different architectures trained in the same way in order to glean insights about the functional role of different structures. Our current work suggests the impact of architecture may be small compared to the impact of training style. It also supports the notion that training only on classification (albeit classification of heavily degraded images) may not be a rich enough task to fully capture the computations the visual system is evolved and developed to perform. This has been argued in the context of purely feedforward CNNs, but it may be especially true for studying the role of recurrence, which is likely more involved in broader aims like scene understanding. While we intentionally studied smaller CNNs trained on a more constrained task in order to make analysis of network behavior and activity tractable, such an approach has limitations. Training recurrent neural networks on more involved tasks might constrain the solution space to better approximate the biological role of recurrence, and also reveal a difference between feedback and lateral recurrence.

That being said, the strategy found by the task-trained networks is not completely unprecedented. Previous work on understanding classification has pointed to the role of auxiliary variables—image features not directly relevant to the classification such as object pose or size—as relevant pieces of information for the network to keep track of (Hong, Yamins, Majaj, & DiCarlo, 2016; Thorat, Aldegheri, & Kietzmann, 2021). The de-noising done by the predictive feedback network results in a loss of information about what noise type was added to the image. The strategy of the task-trained networks, on the other hand, may involve keeping or even enhancing that information and performing classification in light of it rather than despite it.

More generally, this work shows the way in which trained artificial neural networks can be used for hypothesis generation and to demonstrate the broad space of algorithmic solutions to a given problem. For example, previous works have used the presence of de-noising-like dynamics at later visual areas as evidence that recurrence in those areas is important for performance increases (Tang et al., 2014, 2018). We show here that such dynamics in later areas can actually be inherited from very different dynamics at earlier layers—an unexpected mechanism. Analysis of trained neural networks reveals just how counter-intuitively hierarchical and recurrent nonlinear systems like the visual system can behave.

Because of the potential for counter-intuitive mechanisms, large high quality datasets will be crucial for teasing apart the role of recurrence in the visual system. When analyzing our neural networks we had complete access to all units at all time points and in response to all stimuli. This is far from the case for real neural data. And as our analysis of fMRI data shows (Figure 8), real data sources contain much fewer data points, high levels of noise, and large inter-subject variability. The low temporal resolution of fMRI data makes it particularly challenging to use in the study of recurrence. Methods with simultaneously high spatial and temporal resolution that can target multiple brain areas at once (ideally in an expedient way that could allow for many trials and subjects) would provide the best basis for proper analysis of recurrence. So far, nonhuman primate electrophysiology (which provides high spatial and temporal resolution) has struggled with cross-regional recordings. One notable exception is the openly available data from Zandvakili and Kohn (2015), which contains joint V1 and V2 recordings. However, these are in response to very simple oriented gratings while the animal is under anesthesia. Anesthesia is believed to heavily reduce the influence of (particularly feedback) recurrence (Hudetz, 2006), providing a further challenge for collecting the data necessary to study this feature of the visual system.

## 4 Methods

Code for all training and analyses will be posted upon publication.

### 4.1 Network Training

All network training was done with JAX. The architectural parameters of the feedfoward core can be found in Table 1. Weights were initialized with a truncated normal distribution with standard deviation of .02. Five feedforward cores with different random initializations were trained on the standard MNIST dataset using stochastic gradient descent using cross entropy loss for 40000 batches with batch size 256. The weights of the feedforward cores were then held constant as recurrent connections were added.

**Table 1:**
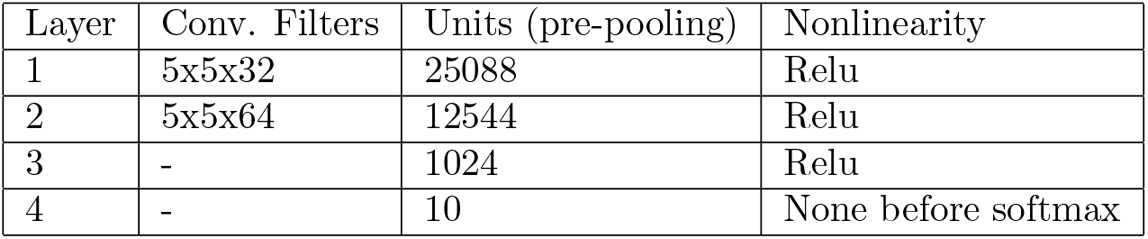
Parameters for feedforward cores

Noise was added to standard MNIST images via four different degradation methods: Gaussian pixel noise, blur, contrast reduction, and occlusion. Noise levels were defined according to the impact they had on the performance of the feedforward core (roughly, level 1 aimed for 90%, 2 for 75%, 3 for 60%, 4 for 45%, 5 for 35% and 6 for 30%). To vary performance levels we varied the intensity of the noise via parameters specific to each noise type. For pixel noise, this was the standard deviation of the distribution that pixel values of the noise mask added to the image were drawn from. For blur, this was the width of the Gaussian blur filter applied to the image. For low contrast, this was the scalar value the pixel intensities were multiplied by (with higher noise meaning lower value). For occlusion, this was the number of 3×3 occlusion blocks (set to the mean pixel intensity of the whole imageset) added to the image.

Task-trained recurrent connections were trained on a dataset made of equal portions of one-fifth standard images and one-fifth of each type of noise-degraded images using the third of five noise levels. For the feedback recurrence networks, 5×5 convolutional connections were added from the 64 feature maps of the second layer to the 32 of the first. For lateral recurrence, 7×7 convolutional connections were added amongst the 32 feature maps of the first layer. In both cases, networks were allowed to run for three total timesteps (a feedforward pass and 2 rounds of recurrence) and losses were calculated using activity at the final timestep. Weights were initialized with a truncated normal distribution with standard deviation of .001 and trained for 40000 batches with batch size 256.

Predictive coding-based feedback was trained according to the procedure described in Choksi et al. (2020). Broadly, the feedback connections were trained using backpropagation on standard (clean) MNIST images to allow the activity at the second layer to predict the activity at the first. This predicted activity is added to the activity of the first layer (parameter values as defined by their method are *α* = 1, λ = .7, *β* = .2). Weights were also initiated with standard deviation .001 and training ran for 40000 batches. The model ran for 13 timesteps, and we sample a step in the middle and the last step for plotting performance over time.

The addition of surround suppression was inspired by Hasani et al. (2019). In that work, a recurrent spatial filter that implements surround suppression within each feature map is fixed into the architecture before training. Here, we simply add such a recurrent filter at the first layer after feedforward training. The weights of this filter were defined by a difference of Gaussians to allow nearby facilitation and surround suppression. We performed hyperparameter tuning on the training set of noisy and clean (standard) MNIST images to obtain the widths of these Gaussians (1.2 and 1.68). This model also ran for three total timesteps.

Two instantiations of each type of recurrence (with the exception of surround suppression which has no random parameters to initialize) were trained for each of the 5 feedforward cores. In all cases, test images used for analysis were distinct from training images.

For comparisons marked ‘grads’ we calculate on an image-by-image basis for the test images the activity changes need to increase classification performance according to the gradients. Specifically, we use backpropagation to calculate the negative partial derivative of layer 1 activity with respect to the cross entropy loss.

### 4.2 Behavioral Analysis

In Figure 2, we compare the classification patterns of the different networks. To compare incorrect label agreement, for each pair of networks we identify images both classify incorrectly. We then report the fraction of those images that are labeled the same way by both networks. To compare images that both networks classify correctly, we analyze the logits (i.e., the activity of the 10-unit final layer). We calculate the logit difference as the logit values at the final time step minus that as the first. We then report the Pearson correlation coefficient for these values, averaged across all images that both networks classify correctly.

### 4.3 Neural Activity Analysis

In Figure 4, we compare the activity changes that occur at the first layer as a result of recurrent influence. To start we take the difference between activity at the final time step minus that at the first and report the Pearson correlation coefficient of these values averaged across all images. To isolate activity changes that may be causally responsible for the performance changes, we first perform principal components analysis on the calculated activity differences. We then reconstruct the activity difference using an increasing number of PCs (ranked by variance explained) and add this difference back onto the feedforward activity. Running this summed activity through the network returns a classification performance for each number of PCs used. Through these we identify the top PCs needed to get the network to 85% of the performance increase (compared to feedforward alone) that is achieved by the full recurrence. We consider these PCs to represent the activity changes that drive the majority of performance changes due to recurrence. To compare across networks, for each pair of networks we report the average principal angles between the subspaces as a measure of how aligned the two subspaces of activity changes are.

To explore the extent to which recurrence ‘de-noises’ the image representation, we created a de-noising metric. First, the correlation between the feedforward activity in response to a clean image and that to its noisy equivalent is calculated. This value is subtracted from the correlation between feedforward activity in response to a clean image and the activity in response to the noisy equivalent after recurrence (i.e., at the final time step):

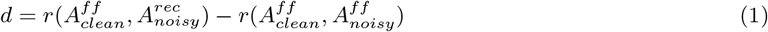

where *r* is the Pearson correlation coefficient and A is activity either before (ff) or after (rec) recurrence in response to the clean or noisy version of an image.

To calculate the dimensionality of neural activity at each layer, the activity was first centered and normalized to unit norm. PCA was performed on the activity (in response to clean and noisy images) separately before and after recurrence. At each time point, the participation ratio of the PCs was calculated according to the formula:

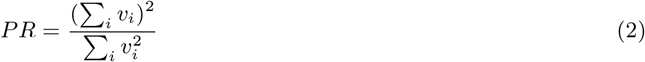

where *v_i_* is the variance explained by the *i^th^* PC.

To estimate the dimensionality of communication across two layers, we first created the covariance matrix of activity across these two layers. Singular value decomposition was then performed on this matrix (note: due to the large number of units in the first and second layers, units were randomly downsampled by 20% for each layer to make this SVD computationally possible). The participation ratio (see above) of these singular values is reported as the cross-layer dimensionality.

### 4.4 fMRI Analysis

We used openly available data associated with the study from Abdelhack and Kamitani (2017, 2018); details of the data collection process can be found there. Briefly, subjects were shown a series of increasingly less blurred versions of 20 natural images until they finally saw the original unblurred version (we use the no-prior conditions of the data wherein subjects were given no information about what to expect in the images). Each image was presented for 8 seconds. Subjects were instructed to guess as to what they believed was in the image and pressed a button to indicate if they were or were not confident in their guess.

For our analysis, we looked at the BOLD response over time for images with the lowest level of blur (as subjects largely were not confident about what they had seen in response to the blurrier images). We normalized each voxel to have 0 mean and unit norm. The time resolution of the BOLD response was 2 seconds, providing 4 time bins per image. We took the first time bin to represent feedforward activity and the last time bin to represent activity after recurrence. We used the same de-noising equation as above to compare the representation of a brain region (composed of a set of voxels) in response to the blurred and clean versions of an image. We also use the same procedure as above to calculate a brain region’s dimensionality.

## Acknowledgements

Many thanks to Mohamed Abdelhack for making the temporal BOLD data from his study available.

## Funding Statement

GWL was funded through a Marie Sklodowska-Curie Individual Fellowship and Sainsbury Wellcome Centre and Gatsby Computational Neuroscience Unit Fellowship.

## Competing interests

The authors have no competing interests to declare.

## Supplementary Figures

**Figure 9:**
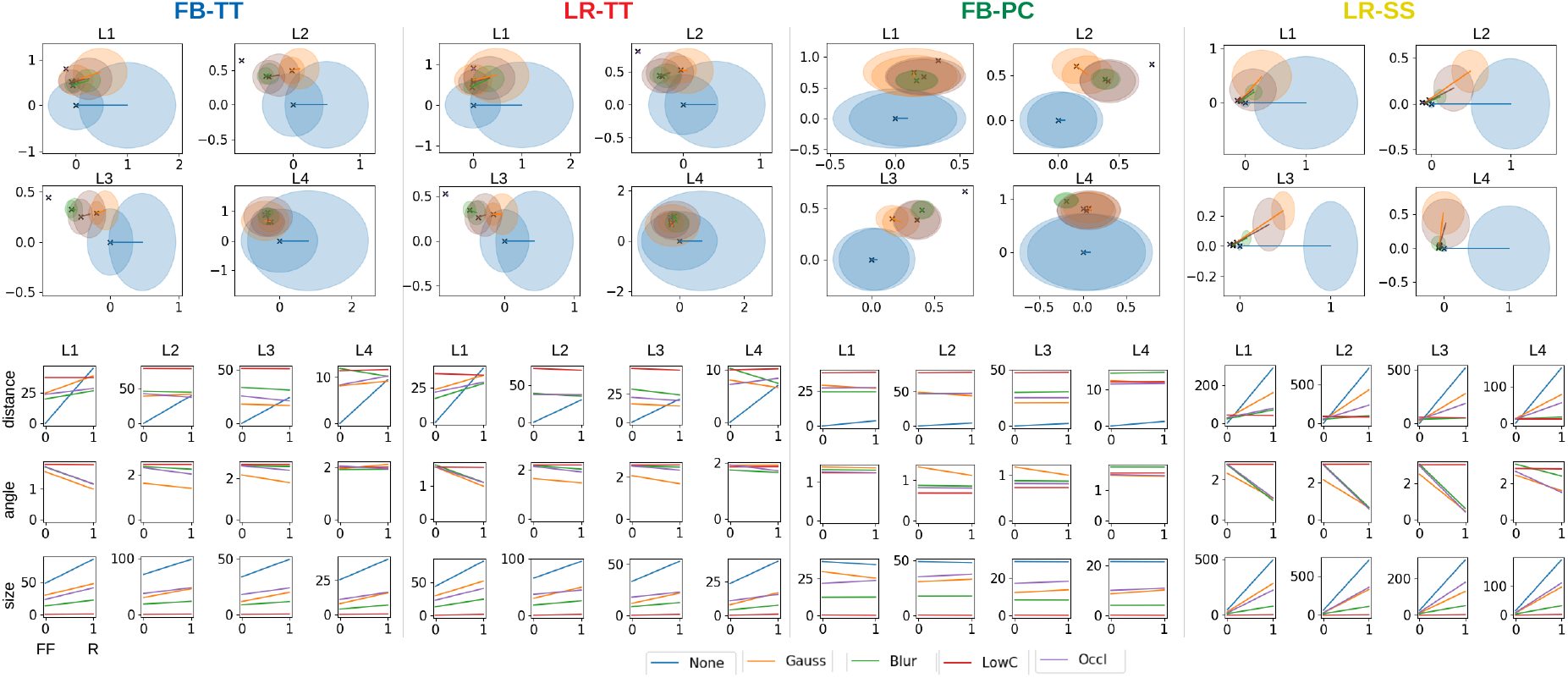
Visualization of how clouds of images move with recurrence. Euclidean distance from the mean feedforward response to clean images and angle from this point are both calculated for means of feedforward responses to each type of noisy image. This is indicated on the top plots by black Xs, with the mean of the clean feedforward response set as the origin and the direction that recurrence moves clean images is set as the x-axis. Diameter of clouds surrounding Xs indicate average Euclidean distance between images of same noise type; color indicates noise type. Lines indicate direction recurrence moves these clouds. Distance, angle, and cloud size values plotted explicitly on bottom.

## Notes

### Competing Interest Statement

The authors have declared no competing interest.

